# Defined neuronal populations drive fatal phenotype in Leigh Syndrome

**DOI:** 10.1101/556019

**Authors:** Irene Bolea, Alejandro Gella, Elisenda Sanz, Patricia Prada-Dacasa, Fabien Menardy, Pablo Machuca-Márquez, Angela Michelle Bard, Franck Kalume, Albert Quintana

## Abstract

Dysfunctions of the mitochondrial energy-generating machinery cause a series of progressive, untreatable and usually fatal diseases collectively known as mitochondrial disease. High energy-requiring organs such as the brain are especially affected, leading to developmental delay, ataxia, respiratory failure, hypotonia, seizures and premature death. While neural affectation is a critical component of the pathology, only discrete neuronal populations are susceptible. However, their molecular identity and their contribution to the disease remain unknown. Mice lacking the mitochondrial Complex I subunit NDUFS4 (Ndufs4KO mice) recapitulate the classical signs of Leigh Syndrome (LS), the most common presentation of mitochondrial disease with predominant CNS affectation. Here, we identify the critical role of two genetically-defined neuronal populations driving the fatal phenotype in Ndufs4KO mice. Selective inactivation of *Ndufs4* in Vglut2-expressing glutamatergic neurons causes brainstem inflammation, motor and respiratory deficits, and early death. On the other hand, *Ndufs4* deletion in GABAergic neurons leads to basal ganglia inflammation without motor or respiratory involvement, but accompanied by severe refractory epileptic seizures preceding premature death. These results provide novel insight in the cell type-specific contribution to LS pathology and open new avenues to understand the underlying cellular mechanisms of mitochondrial disease.

## Introduction

Leigh syndrome (LS) is the most frequent pediatric mitochondrial disorder, leading to defective mitochondrial energy metabolism. LS affects 1 in 40,000 births^1^, although adult onset has also been described^2^. Mutations in more than 75 genes have been described to cause LS^3^. To date, no effective treatment or cure exists. Clinically, albeit highly variable, LS symptoms usually include failure to thrive, hypotonia, rigidity, seizures, ataxia, lactic acidosis, encephalopathy and premature death^1,3,4^. LS is characterized by restricted anatomical and cellular specificity^5^, a common feature shared by mitochondrial diseases, affecting high energy-requiring tissues such as muscle and brain^6^. Pathologically, LS is characterized by the presence of bilateral symmetrical lesions predominantly in the brainstem and basal ganglia^5^. Neuronal damage is responsible for most of the fatal symptoms, including respiratory failure and seizures^7^. However, the identity of the affected cellular populations and the molecular determinants of neuronal vulnerability have not been adequately elucidated, representing a challenge for the development of efficient treatments.

Mutations affecting the NDUFS4 subunit of mitochondrial Complex I, a key structural component for the assembly, stability and activity of the complex^8^, are commonly associated with a severe, early-onset LS phenotype^9^. Although a late-onset case of LS has been recently reported^10^, prognosis is usually poor and most of the patients die in early childhood^4^.

Animals with a global deletion of *Ndufs4* (Ndufs4KO mice) develop a fatal encephalomyopathy which recapitulates the classical signs of LS, including motor alterations, respiratory deficits, epilepsy and premature death^11,12^. Behavioral and neuropathological characterization of Ndufs4KO mice revealed the pivotal role of the dorsal brainstem, particularly the vestibular nucleus (VN), in disease manifestation and progression^13^. However, the genetic identity of the neuronal populations and circuitries involved in the plethora of symptoms observed have not yet been identified.

Here, we describe the contribution of genetically defined, discrete neuronal populations to the fatal phenotype of Ndufs4KO mice. To that end, we generated three mouse lines using a conditional genetic approach that selectively inactivates *Ndufs4* in glutamatergic (Vglut2-expressing), GABAergic (Gad2-expressing) or cholinergic (ChAT-expressing) neurons. The results reveal distinct, lethal phenotypes for the glutamatergic and GABAergic neuronal populations.

## Results

### Reduced lifespan and body weight in cell type-specific conditional Ndufs4KO mice

To dissect the neuronal cell types contributing to the neuropathology observed in Ndufs4KO mice^11–13^, we generated three mouse lines lacking *Ndufs4* selectively in glutamatergic (expressing Vglut2), GABAergic or cholinergic neurons. We did this by crossing *Ndufs4* exon2-floxed mice with Cre-driver lines of mice expressing either *Slc17a6-Cre, Gad2-Cre* or *Chat-Cre* as described in Figure 1A and Supplementary Figure S1A; the affected mice are referred to here as: Vglut2:Ndufs4cKO, Gad2:Ndufs4cKO or ChAT:Ndufs4cKO mice and their respective controls are Vglut2:Ndufs4cCT, Gad2:Ndufs4cCT or ChAT:Ndufs4cCT.

**Figure 1.**
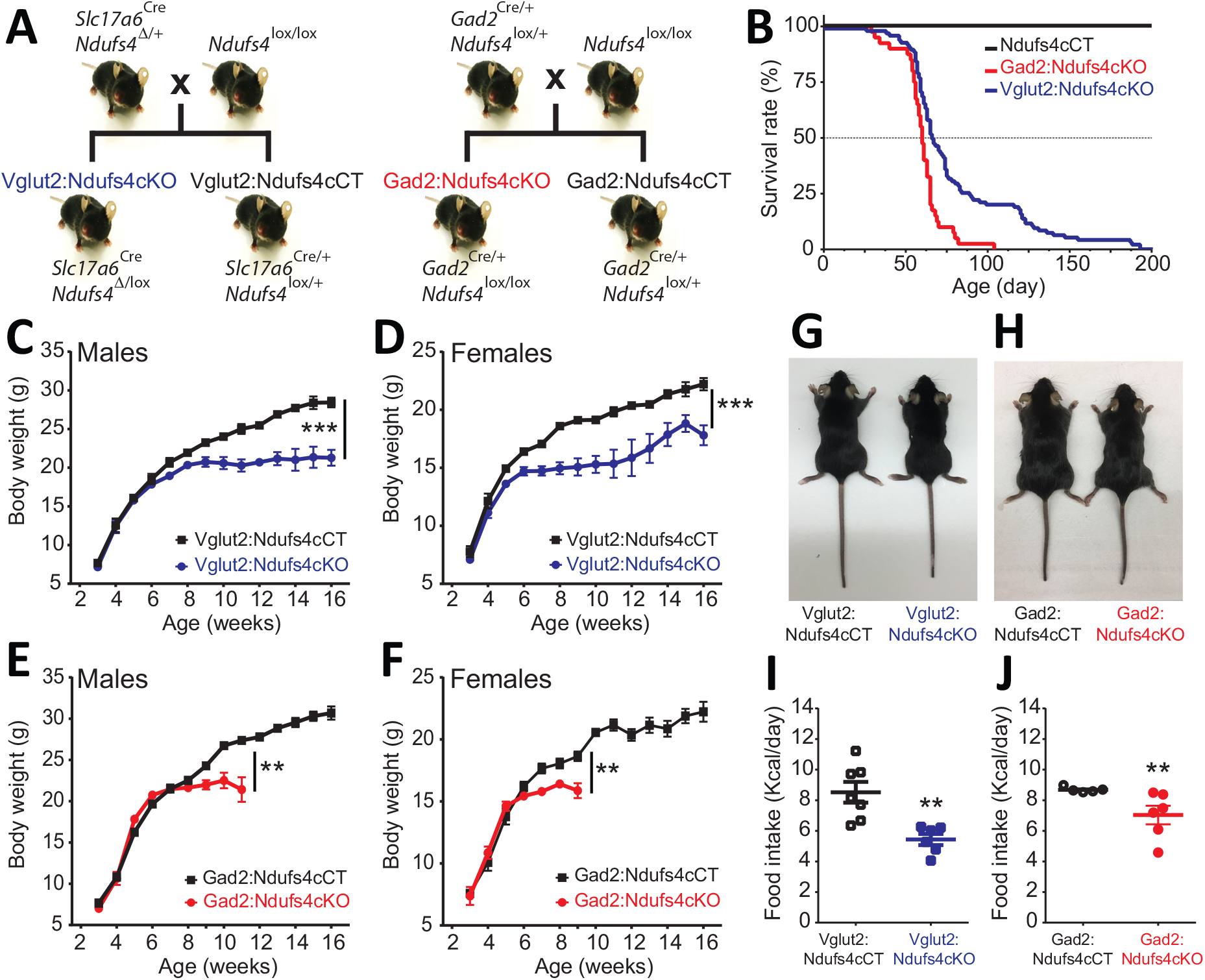
Generation of two mouse lines lacking *Ndufs4* selectively in Vglut2-expressing glutamatergic neurons or Gad2-expressing GABAergic neurons. (A) Breeding strategy to obtain mice with conditional *Ndufs4* deletion in glutamatergic neurons (Vglut2:Ndufs4cKO) or GABAergic neurons (Gad2:Ndufs4cKO) and their respective controls (Vglut2:Ndufs4cCT, Gad2:Ndufs4cCT). (B) Kaplan-Meier survival curve for Vglut2:Ndufs4cKO (n=96, blue), Gad2:Ndufs4cKO (n=40, red) and cCT mice (n=50, black). (C-D) Body weight curves for Vglut2:Ndufs4cKO (blue, n=45 males, n=40 females) and Vglut2:Ndufs4cCT (black, n=35 males, n=40 females). (E-F) Body weight curves for Gad2:Ndufs4cKO (red, n=14 males, n=19 females) and Gad2:Ndufs4cCT (black, n=25 males, n=35 females). Data are presented as the mean ± SEM. Statistical analysis was performed using a two-way ANOVA followed by Bonferroni post-test: \*\**P* < 0.01, \*\*\**P* < 0.001. (G) Representative images showing reduced body size in Vglut2:Ndufs4cKO mice when compared to Vglut2:Ndufs4cCT mice at P65. (H) Gad2:Ndufs4cKO mice also display reduced body size when compared to control littermates (Gad2:Ndufs4cCT) at P70. Food intake values (kcal/day) for Vglut2:Ndufs4cKO (n=6, closed blue squares) and Vglut2:Ndufs4cCT (n=7, open black squares) mice (I) and Gad2:Ndufs4cKO (n=6, closed red circles) and Gad2:Ndufs4cCT (n=5, open black circles) mice (J) at 8 weeks of age. Data are presented as the mean ± SEM. \*\**P* < 0.01. Statistical analysis was performed using an unpaired *t-* test (I) or a Mann Whitney U test (J).

*Ndusf4* gene inactivation in glutamatergic or GABAergic neurons of both male and female mice resulted in failure to thrive and premature death (Figure 1B-F); however, there was no effect on survival, body weight, or motor function when *Ndufs4* expression was abolished in cholinergic neurons (Supplemental Figures S1B-C). Selective deletion of *Ndufs4* in glutamatergic or GABAergic neurons was confirmed by western blot analysis of NDUFS4 levels in brain areas where Vglut2 or Gad2 are preferentially expressed^14^ (Supplemental Figure S2A). Vglut2:Ndufs4cKO mice had a median lifespan of 67 days with a mortality rate of 90% at postnatal day (P) 128. Similarly, Gad2:Ndufs4cKO mice had a median lifespan of 60 days (90% mortality at P70) (Figure 1B). This premature death was preceded by weight loss in male and female mice of both genotypes (Figure 1C-F). At 7 weeks of age for females and 9 weeks of age for males, Vglut2:Ndufs4cKO mice stopped gaining weight, which resulted in an overall reduction in body weight when compared to age-matched controls (Figure 1C-D). Similarly, male and female Gad2:Ndufs4cKO mice started to show reduced body weight 2–3 weeks before manifesting a sudden unexpected death (Figure 1E-F). Both Vglut2:Ndufs4cKO and Gad2:Ndufs4cKO mice were also significantly smaller than their littermate controls (Figure 1G-H). This lack of weight gain and reduced size appeared to be due to decreased food intake in both genotypes (Figure 1I-J); however, this was not significantly different when food consumption was normalized to body weight (Figure S2B).

### Vglut2:Ndufs4cKO and Gad2:Ndufs4cKO mice manifest clinically distinct phenotypes

The reduced lifespan and decreased body weight observed in both Vglut2:Ndufs4cKO and Gad2:Ndufs4cKO mice were the result of two prominently different clinical presentations. Gad2:Ndufs4cKO mice were, for the most part, phenotypically indistinguishable from controls, without any overt clinical alteration beyond a reduced growth rate for a few weeks prior to a sudden premature death. On the other hand, Vglut2:Ndufs4cKO mice manifested progressive motor and respiratory deterioration with most of the clinical signs visibly apparent (Table 1).

**Table 1.**
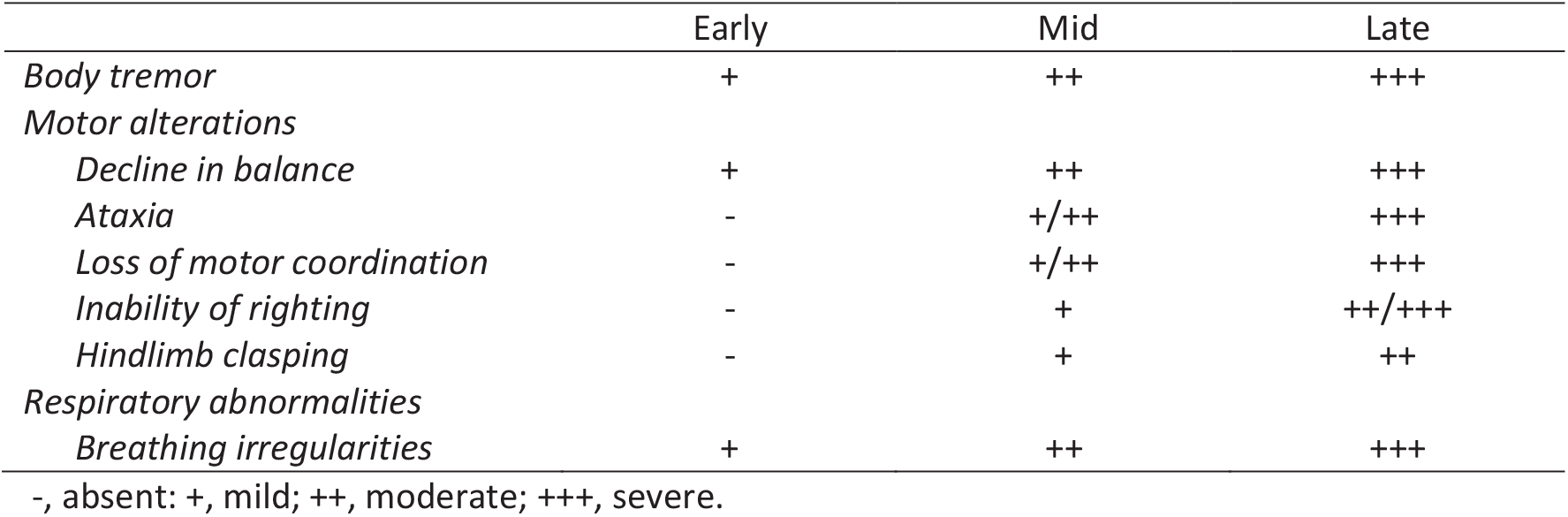
Neurological signs observed in Vglut2:Ndufs4cKO mice according to the stage of disease.

Vglut2:Ndufs4cKO mice presented body tremor and a decline in balance as early as at 5 weeks (early stage of the disease), which dramatically worsened as the disease progressed. In a mid-stage of the disease, mice increased body tremor and showed a prominent decline in balance and motor coordination. Subsequently, animals started exhibiting ataxia and a progressive loss of the righting and hindlimb extension reflexes (Supplemental Figure S3A). At a late stage of the disease, these mice showed increased tremor, were completely docile and hypotonic, and would not explore new environments. Animals also had difficulty maintaining a regular breathing pattern at an early stage of the disease. These breathing abnormalities worsened as the disease progressed with mice presenting noticeably shorter and deeper respirations at advanced stages of the disease (Supplemental Video S1). Furthermore, at a late stage of the disease, about 40% of Vglut2:Ndufs4cKO developed hindlimb dragging and eventually hindlimb paralysis. We observed increased glial reactivity but no neuronal loss in the spinal cord of these mice when compared to Vglut2:Ndufs4cCT mice, which was determined by immunoblot analysis of spinal cord lysates from Vglut2:Ndufs4cCT and Vglut2:Ndufs4cKO mice using antibodies against GFAP (glial fibrillary acidic protein), Iba-1 (ionized calcium-binding adapter molecule 1) and NSE (neuronal specific enolase) (Supplemental Figure S3B). This glial reactivity was further confirmed by immunofluorescence analysis using anti-GFAP and anti-Iba-1 antibodies in the spinal cords of the Vglut2:Ndufs4cKO mice that exhibited hindlimb dragging (Supplemental Figure S3C, top panels). However, no signs of demyelination or immune cell infiltration were observed (Supplemental Figure S3C, mid and bottom panels).

### Neuronal identity defines distinct neuroinflammatory patterns after *Ndufs4* deletion

LS is characterized by symmetrical brain lesions and neuroinflammation in select nuclei, predominantly brainstem and/or basal ganglia^5^. Accordingly, Ndufs4-deficient mice present overt lesions in brainstem (namely vestibular nucleus (VN), cerebellar fastigial nucleus (FN) and inferior olive (IO)), olfactory bulb and basal ganglia^11,13,15^. To define the contribution of *Ndufs4* deficiency in either excitatory or inhibitory neurons to the overall neuroinflammatory phenotype and identify the specific brain areas with exacerbated astroglial and microglial reactivity, we performed immunofluorescence analysis using anti-GFAP and anti-Iba1 antibodies on brain sections of Vglut2:Ndufs4cKO and Gad2:Ndufs4cKO mice, and their respective controls (Figures 2A-B). Analysis of these sections showed marked glial reactivity in VN, IO and FN (Figure 2A) in Vglut2:Ndufs4cKO mice, recapitulating most of the neuroinflammatory profile described for the global Ndufs4KO mice^11^. In contrast, Gad2:Ndufs4cKO mice presented a more restricted glial reactivity pattern, including marked microglial and astroglial reactivity in primarily GABAergic nuclei such as the external globus pallidus (GPe) in the basal ganglia and the substantia nigra pars reticulata (SNr) (Figure 2B), without affecting neighboring non-GABAergic areas like the dopaminergic substantia nigra pars compacta (SNc) (Supplemental Figure S4A). Other prominently GABAergic areas such as the olfactory bulb also showed increased immunoreactivity for GFAP and Iba-1 in Gad2:Ndufs4cKO mice when compared to control mice (Figure 2B). In a percentage of Gad2:Ndufs4cKO mice, prominent microglial reactivity with Purkinje neuron loss (as assessed by calbindin staining) was also observed in the cerebellar vermis and flocculus (Supplemental Figure S4B).

**Figure 2.**
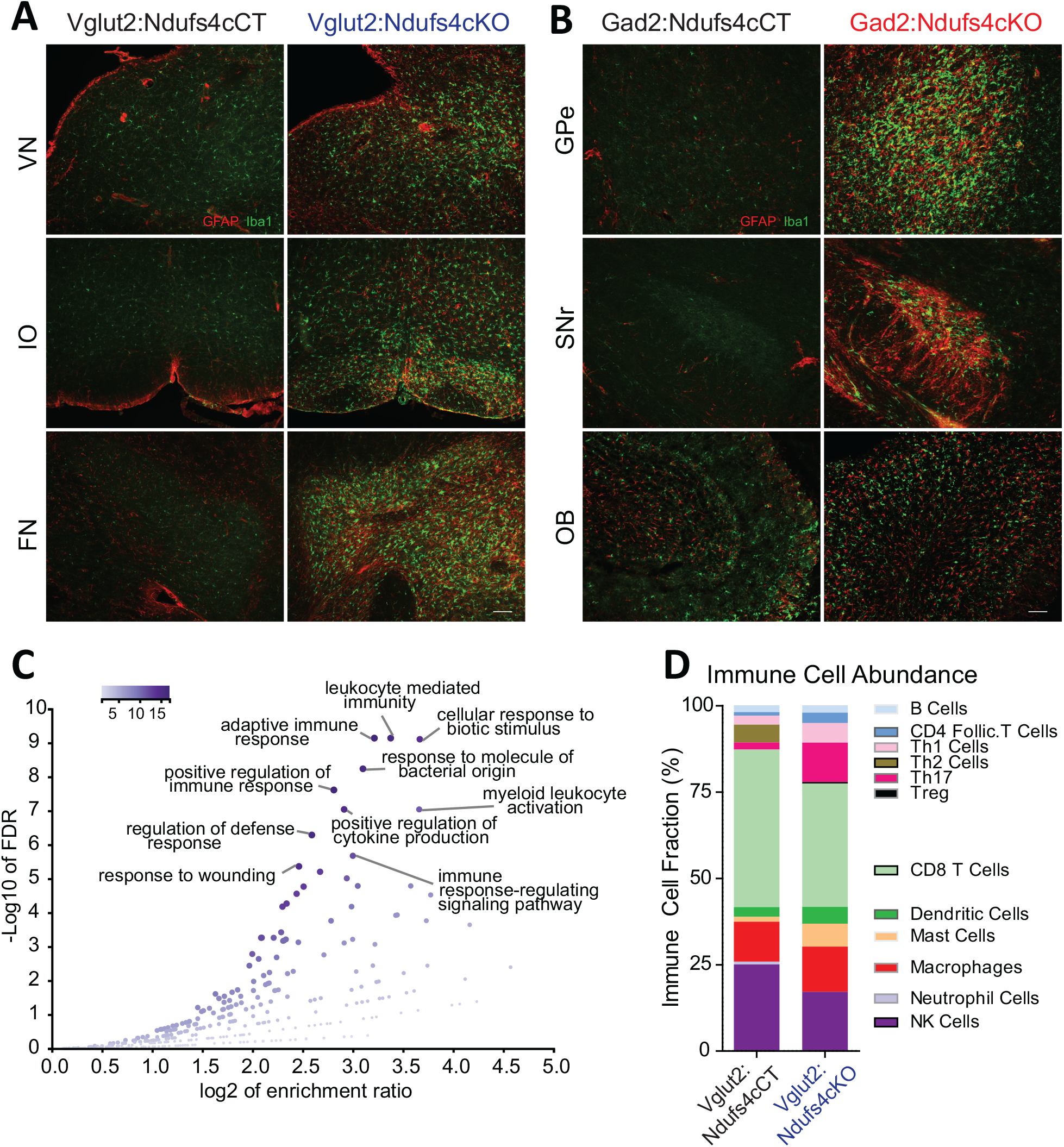
Distinct histopathological pattern in Vglut2:Ndufs4cKO and Gad2:Ndufs4cKO mice. (A) Micrographs showing GFAP (red) and Iba-1 (green) staining in the vestibular nucleus (VN), inferior olive (IO) and the fastigial nucleus (FN) of Vglut2:Ndufs4cKO mice and their respective controls (Vglut2:Ndufs4cCT). (B) Micrographs showing GFAP and Iba-1 staining in the external globus pallidus (GPe), the sustantia nigra pars reticulata (SNr) and the olfactory bulb (OB) in Gad2:Ndufs4cKO and Gad2:Ndufs4cCT mice. (C) Volcano plot showing the top 10 most enriched Biological Process (non-redundant) categories by Gene Ontology (GO) Overrepresentation Analysis (ORA) from differentially expressed genes (1.4-fold or higher) in the brainstem of Vglut2:Ndufs4cKO mice (n=4) when compared to Vglut2:Ndufs4cCT mice (n=4). (D) ImmuCC analysis showing the relative immune cell abundance inferred from gene expression analysis in the brainstem of Vglut2:Ndufs4cKO and Vglut2:Ndufs4cCT mice.

The extensive neuroinflammation present in the brainstem of Vglut2:Ndufs4cKO mice allowed us to further define this inflammatory phenotype using whole-tissue transcriptional profiling. Gene expression analysis in the brainstem of Vglut2:Ndufs4cKO mice using Illumina Beadchips (MouseRef-8 V2; Illumina) showed that differentially expressed (DE) mRNAs were, for the most part, upregulated in Vglut2:Ndufs4cKO mice (*P*<0.05, Fold Change>2) (Supplemental Figure S4C). Selected genes included transcripts directly involved in the regulation of the immune system such as chemokines and their receptors (*Ccl3, Ccl4* and *Cxcr3*), toll-like receptors *(Tlr2, Tlr7),* complement proteins (*C1qa, C1qb, C3* and *C4b*), surface antigens (*Cd84, Cd86* and *Ly9*), and markers of the myeloid cell lineage (infiltrating macrophages and microglia) including *Lyz, Lyz2, Lyzs* and *Slc11a1*, among others.

Functional enrichment analysis of differentially expressed mRNAs (1.4-fold or higher) using overrepresentation analysis (ORA)^16^ showed that the top 10 most overrepresented Gene Ontology (GO) categories (biological process, non-redundant) were all related to defense and immune responses, and also uncovered components of the adaptive immune response (GO:0002250; *P*=2.0177e-12, FDR=7.2033e-10) including “leucocyte mediated immunity” (GO:0002443; *P*=1.7533e-12, FDR=7.2033e-10) and myeloid leukocyte activation (GO:0002274; *P*= 8.7458e-10, FDR=9.0537e-8) (Fig. 2C). Therefore, to further characterize the immune cell composition in these whole tissue gene expression profiles, we applied recently developed deconvolution tools that use leucocyte gene expression signature matrices to computationally infer the relative proportions of each immune cell type in gene-expression mixtures^17,18^ (Figure 2D). This analysis revealed a gene expression profile consistent with an increased proportion of proinflammatory CD4 cells (Follicular cells, Th1, Th17 and Treg), dendritic cells, mast cells and macrophages, and an underrepresentation of CD4 Th2 cells, CD8 cells and NK cells in the brainstem of Vglut2:Ndufs4cKO mice compared to controls, showing that *Ndufs4* deficiency in glutamatergic neurons promotes a neuroinflammatory environment that involves a commitment to distinct proinflammatory TH cell lineages and a defined profile of tissue defense cells.

### Vglut2:Ndufs4cKO but not Gad2:Ndufs4cKO mice present motor alterations and respiratory deficits

Motor dysfunction is a prominent feature in LS and Ndufs4KO mice pathology^4,11^. To genetically define the neuronal cell types mediating this functional disorder, we assessed motor coordination in Vglut2:Ndufs4cKO mice, Gad2:Ndufs4cKO mice, and their respective controls (Figure 3). Starting at P40, and concurring with the onset of clinical signs, Vglut2:Ndufs4cKO mice showed impaired rotarod performance when compared to control littermates (Figure 3A). While control mice maintained rotarod performance, Vglut2:Ndufs4cKO mice failed to properly execute the task, presenting a progressive decline in motor coordination, in line with the clinical phenotype. Conversely, and in agreement with the lack of apparent clinical signs, no differences in rotarod performance were observed in Gad2:Ndufs4cKO mice compared to control littermates (Figure 3B). Exposure to a novel environment revealed a hypoactive phenotype in Vglut2:Ndufs4cKO mice as assessed by a reduction in the total distance traveled and the speed of exploratory movement in the open-field test (Figures 3C-E). The severity of this phenotype in Vglut2:Ndufs4cKO mice positively correlated with age and disease stage (Supplemental Figure S5). In contrast, no significant differences were observed in either distance traveled or speed in the open-field test between Gad2:Ndufs4cKO mice and their respective controls (Figures 3F-H).

**Figure 3.**
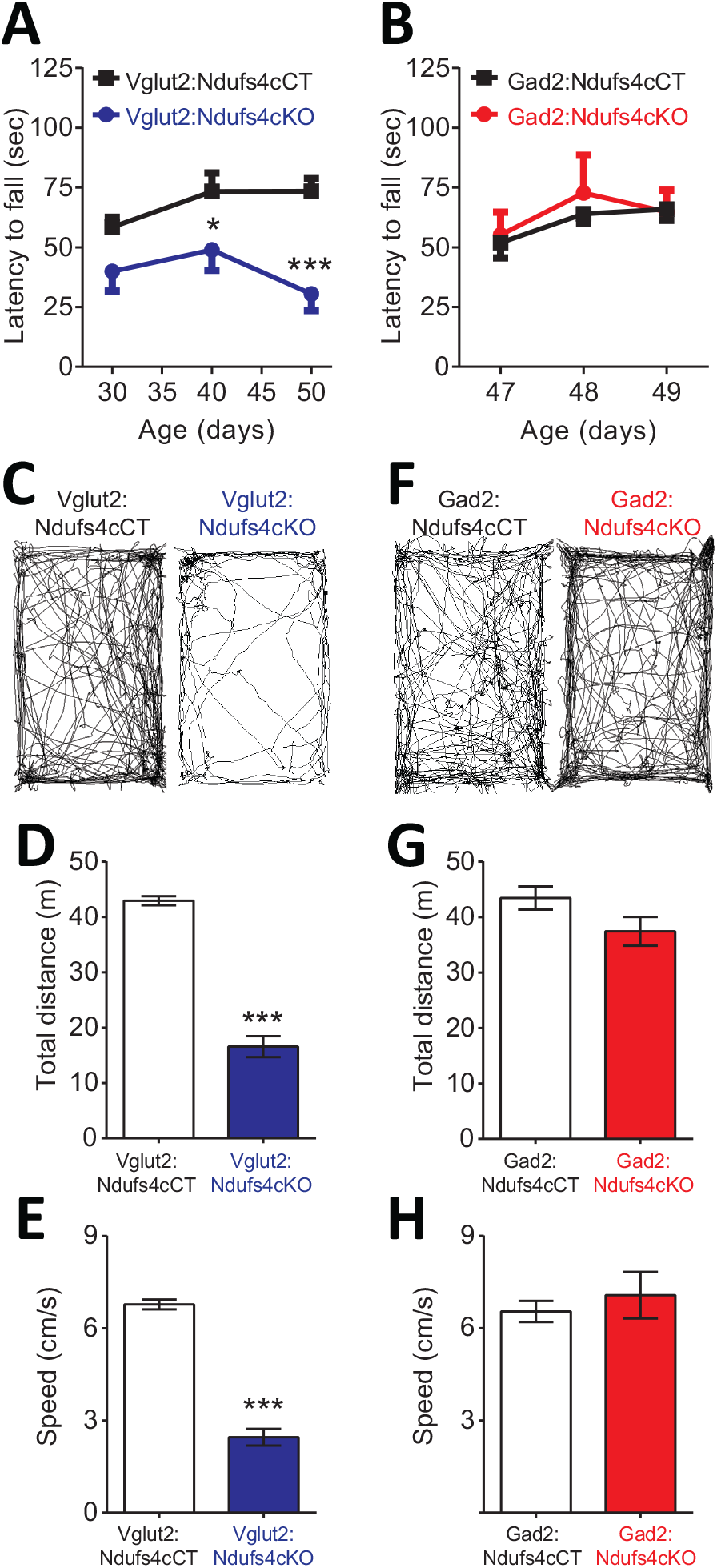
Motor impairment in Vglut2:Ndufs4cKO mice. Latency to fall (seconds) in Rotarod test for Vglut2:Ndufs4cKO (n=8) and Vglut2:Ndufs4cCT mice (n=10) (A), and Gad2:Ndufs4cKO (n=4) and Gad2:Ndufs4cCT mice (n=10) (B). Representative mouse track plots, total distance traveled and average speed during 10 minutes in the open filed for Vglut2:Ndufs4cKO (n=11) and Vglut2:Ndufs4cCT mice (n=12) (C-E), and Gad2:Ndufs4cKO (n = 12) and Gad2:Ndufs4cCT mice (n= 8) (F-H). Data are presented as the mean ± SEM. Statistical analysis was performed using one-way ANOVA followed by Bonferroni post-test (\**P* < 0.05, \*\*\**P* < 0.001).

Respiratory abnormalities are frequently associated with disease mortality in LS patients and global Ndufs4KO mice^5,13,19^. To provide insight into the genetic identity of the neurons responsible for this respiratory phenotype, we assessed respiratory function by unrestrained whole-body plethysmography in awake Vglut2:Ndufs4cKO and Gad2:Ndufs4cKO mice. Vglut2:Ndufs4cKO mice exhibited erratic plethysmographic recordings (Figure 4A). In these mice, the frequency of respiration (f_R_) was markedly reduced at a mid-stage of the disease and worsened as disease progressed (Figure 4C). In addition, significant differences were also seen in the volume of air inspired by the animal during one breath (tidal volume, V_T_). V_T_ was increased in Vglut2:Ndufs4cKO mice at a mid-stage of the disease and became significantly larger than in Vglut2:Ndufs4cCT at late disease stages (Figure 4D). In contrast, Gad2:Ndufs4cKO mice did not exhibit irregular plethysmographic traces and had breathing patterns (f_R_ and V_T_) that did not differ from controls (Figures 4B and 4E-F). These data reveal that the motor impairment and respiratory deficits observed in the Ndufs4KO mouse are mediated by Vglut2-expressing excitatory neuronal populations.

**Figure 4.**
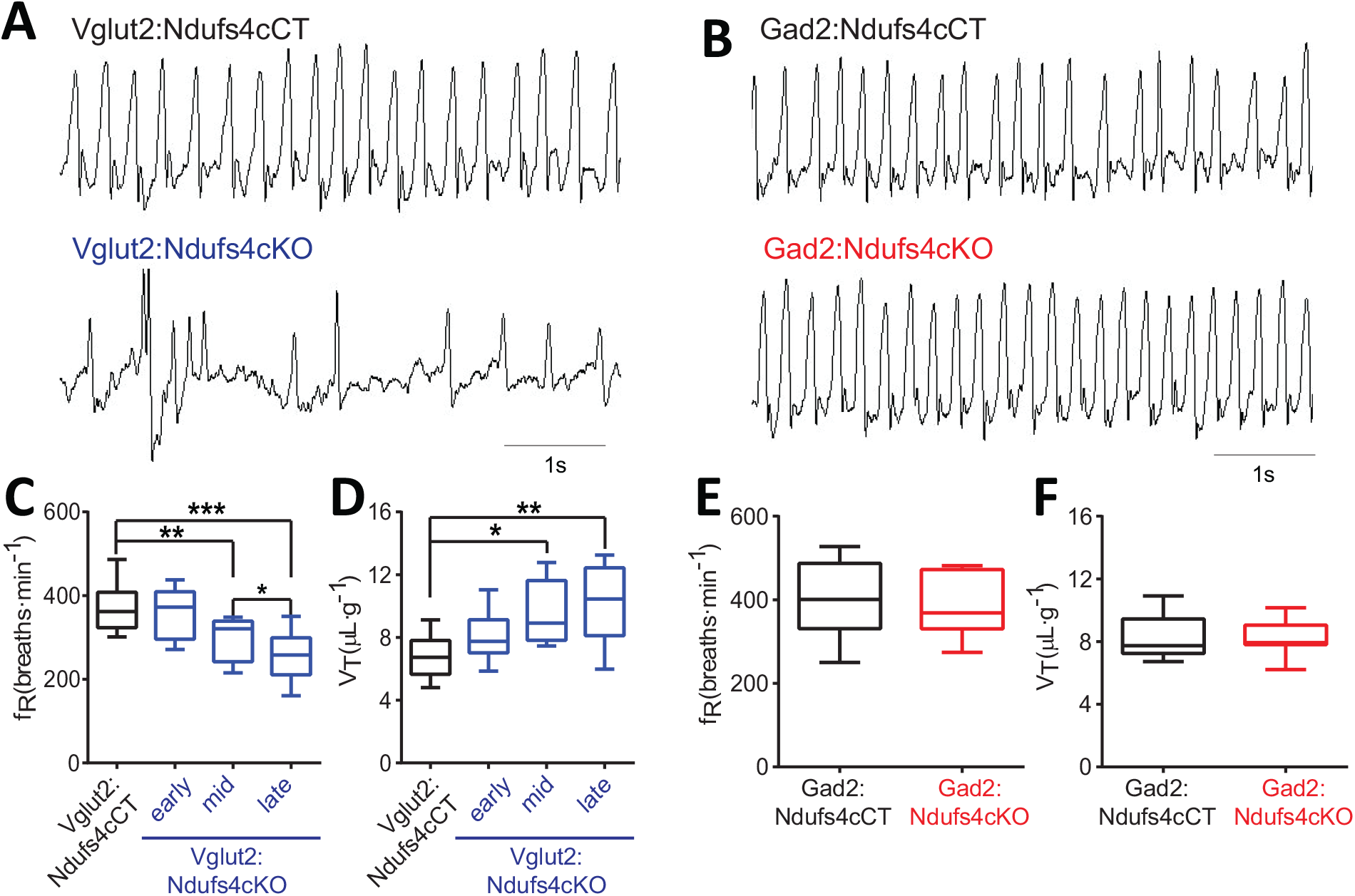
Breathing alterations in Vglut2:Ndufs4cKO mice. Representative 5-second plethysmographic recordings from (A) Vglut2:Ndufs4cCT mice (top), late stage Vglut2:Ndufs4cKO mice (bottom), and (B) Gad2:Ndufs4cCT mice (top) and Gad2:Ndufs4cKO (bottom). Respiratory frequency (fR, breaths-min^-1^) and tidal volume (V_T_, μL-g^-1^) for Vglut2:Ndufs4cCT (n=12) and Vglut2:Ndufs4cKO mice at different stages of the disease (early, n=7; mid, n=8; late, n=12) (C-D) and Gad2:Ndufs4cCT (n=7) and Gad2:Ndufs4cKO (n=7) mice (E-F). Data are presented as the mean ± SEM. Statistical analysis was performed using one-way ANOVA followed by Bonferroni post-test (*P < 0.05, \*\**P* < 0.01, \*\*\**P* < 0.001).

We have shown that neuronal inactivation of *Ndufs4* in the VN promotes breathing abnormalities^13^. This observation, along with the extensive neuroinflammation observed in the VN of Vglut2:Ndufs4cKO mice, prompted us to assess the activity of Vglut2-expressing neurons in the VN using *in vivo* electrophysiology. Cell-firing activity was recorded during an open-field session before identification by laser stimulation of Channelrhodopsin-2 (ChR2)-expressing vestibular glutamatergic neurons in Vglut2:Ndufs4cKO and Vglut2:Ndufs4cCT mice (Figure 5A). Electrophysiological recordings *in vivo* showed that the firing rate for vestibular Vglut2-expressing neurons from Vglut2:Ndufs4cKO mice was not significantly different than controls when mice were at rest. However, when the mice were actively moving in the open field, the Vglut2-expressing VN neurons in the control mice approximately doubled their firing rate to about 40 Hz but the experimental group did not (Figure 5B). The failure of VN glutamatergic neurons in Vglut2:Ndufs4cKO mice to respond to motor activity may account for the breathing abnormalities.

**Figure 5.**
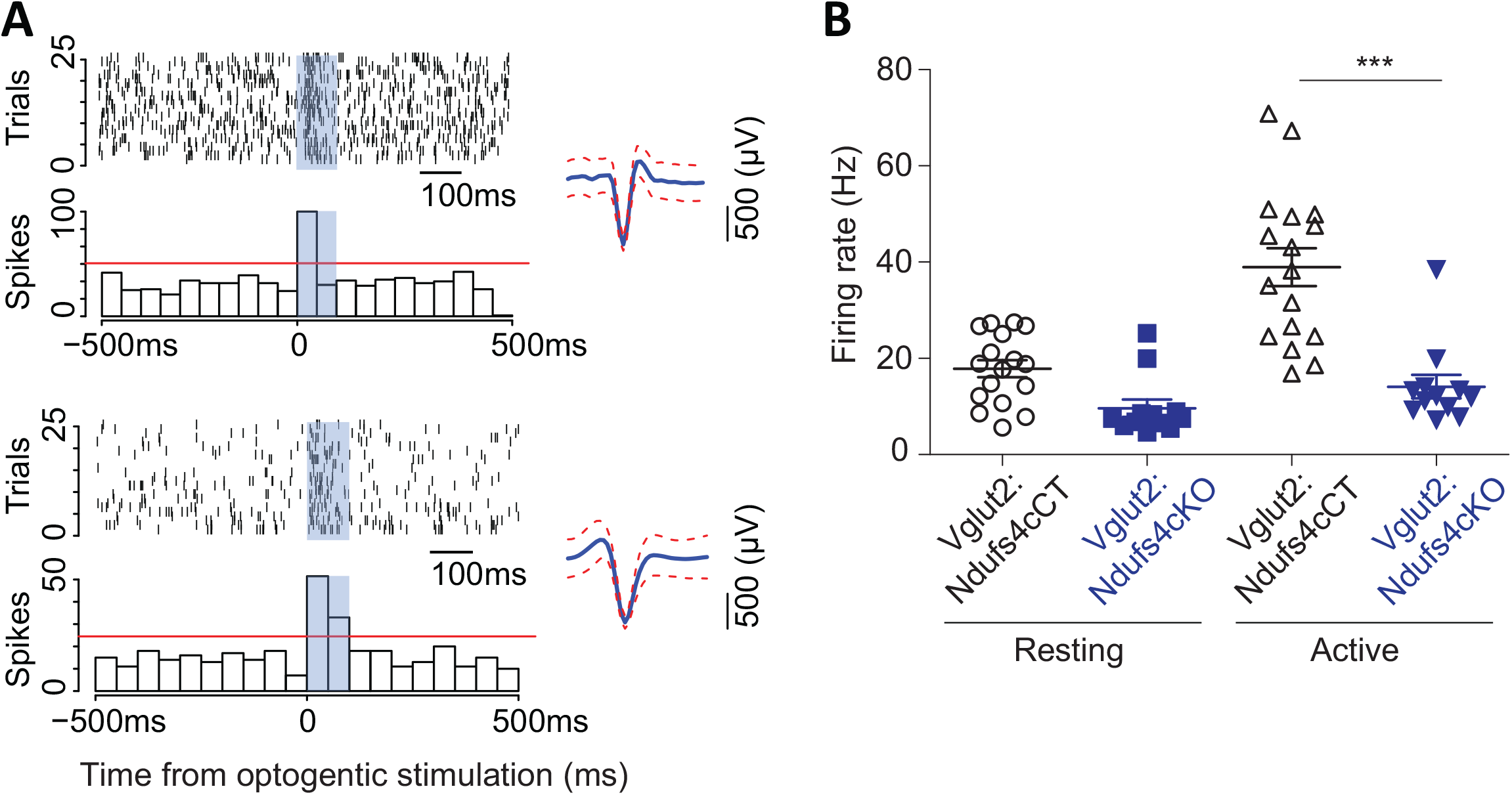
Vglut2-expressing glutamatergic cells in the vestibular nucleus show reduced *in vivo* electrophysiological activity in Vglut2:Ndufs4cKO mice. (A) Representative identification of two different glutamatergic cells with raster plots (upper part) and peri-stimulus histograms (PSTH, lower part) using optogenetic stimulation. 0 represents the onset of optogenetic stimulation. Red line represents mean firing rate from the 500ms before the stimulus plus 3 time the SD. Blue shading indicates period of laser stimulation. (B) Firing rate of Vglut2-expressing glutamatergic cells during a 10-min session in an open-field. Reduced electrophysiological activity was observed in freely-moving Vglut2:Ndufs4cKO mice when compared to Vglut2:Ndufs4cCT mice regardless behavioral state (resting or active). Data are presented as the mean ± SEM. Statistical analysis was performed using two-way repeated measures ANOVA with Bonferroni post-test (\*\*\**P* < 0.001).

### Gad2:Ndufs4cKO manifest sudden unexpected death associated with epilepsy

In Gad2:Ndufs4cKO mice, spontaneous seizure-like events were observed during routine husbandry practices such as lifting the animal by the tail or cage cleaning (Supplemental Video S2). To define whether fatal seizures were the cause of the unanticipated death observed in Gad2:Ndufs4cKO mice, starting at P40 animals were continuously video-recorded to monitor for potential seizures leading to an abrupt death. Analysis of the recordings revealed that all deaths in Gad2:Ndufs4cKO mice consistently followed a severe generalized tonic-clonic convulsion. In contrast, spontaneous, sporadic or lethal seizures were not observed in either controls (data not shown) or Vglut2:Ndufs4cKO mice (Supplemental Figure S6A-C). Detailed visualization and analysis of the images unveiled that seizures in Gad2:Ndufs4cKO mice were mostly generalized and of multiple semiology (unilateral, primary bilateral, secondary bilateral or alternating). Subsequent electroencephalographic (EEG) and electromyographic (EMG) characterization showed the presence of spontaneous generalized tonic-clonic (GTC) seizures (Figure 6A), interictal spikes and myoclonic seizures in Gad2:Ndufs4cKO mice (Figure 6B-C), which are hallmarks of epilepsy in mitochondrial disorders^20^, with 40-60% of mitochondrial disease patients manifesting seizures^21–23^. In addition, local field potential (LFP) recordings identified the presence of seizure-like events^24^ in the GPe of Gad2:Ndufs4cKO mice as soon as P35, while being absent in control mice (Figure 6D), suggesting that subcortical alterations in affected regions may precede the development of generalized seizures.

**Figure 6.**
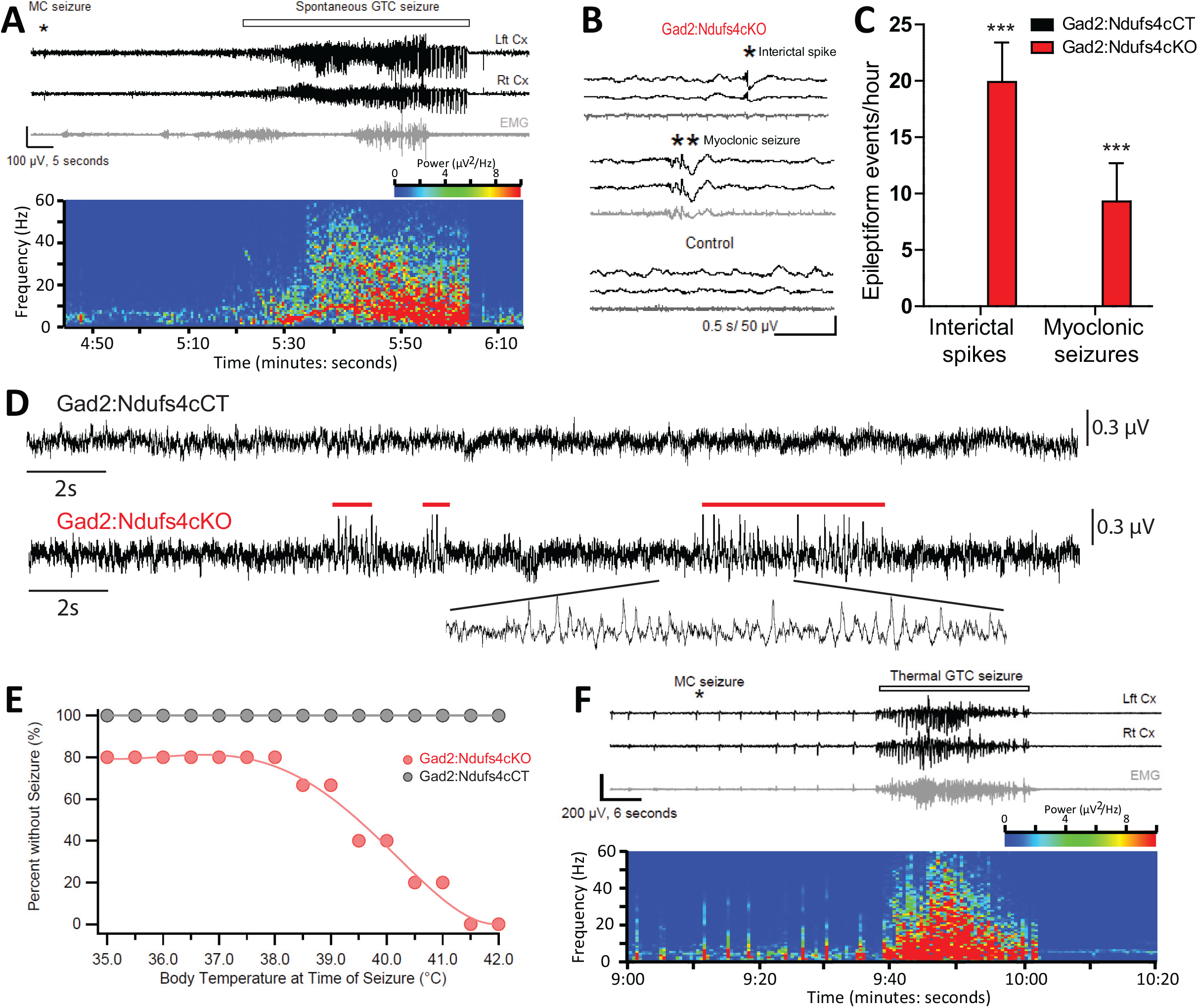
Gad2:Ndufs4cKO mice present epileptic seizures. (A) EEG-EMG recordings and spectrogram analysis showing the frequency and power density of a typical spontaneous seizure in a Gad2:Ndufs4cKO mouse. A preceding myoclonic seizure (MC) is marked with an asterisk. Representative EEG-EMG traces (B) and number of epileptiform events (interictal spikes and myoclonic seizures) (C) in Gad2:Ndufs4cKO and control (Gad2:Ndufs4cCT) mice. Data are presented as the mean ± SEM. Statistical analysis was performed using two-way ANOVA (\*\*\**P* < 0.001). (D) Local field potential (LFP) recordings in the globus pallidus of Gad2:Ndufs4cCT and Gad2:Ndufs4cKO mice at P35. Notice the presence of epileptic events (red lines) in the traces from Gad2:Ndufs4cKO mice. (E) Percentage of Gad2:Ndufs4cKO and Gad2:Ndufs4cCT mice remaining seizure-free after increasing body temperature. (F) Representative EEG-EMG recordings and spectrogram analysis of a thermally-induced seizure in a Gad2:Ndufs4cKO mouse. A preceding myoclonic seizure (MC) is marked with an asterisk. Lft Cx, left cortex; Rt Cx: right cortex; EMG: electromyography.

**Figure 7.**
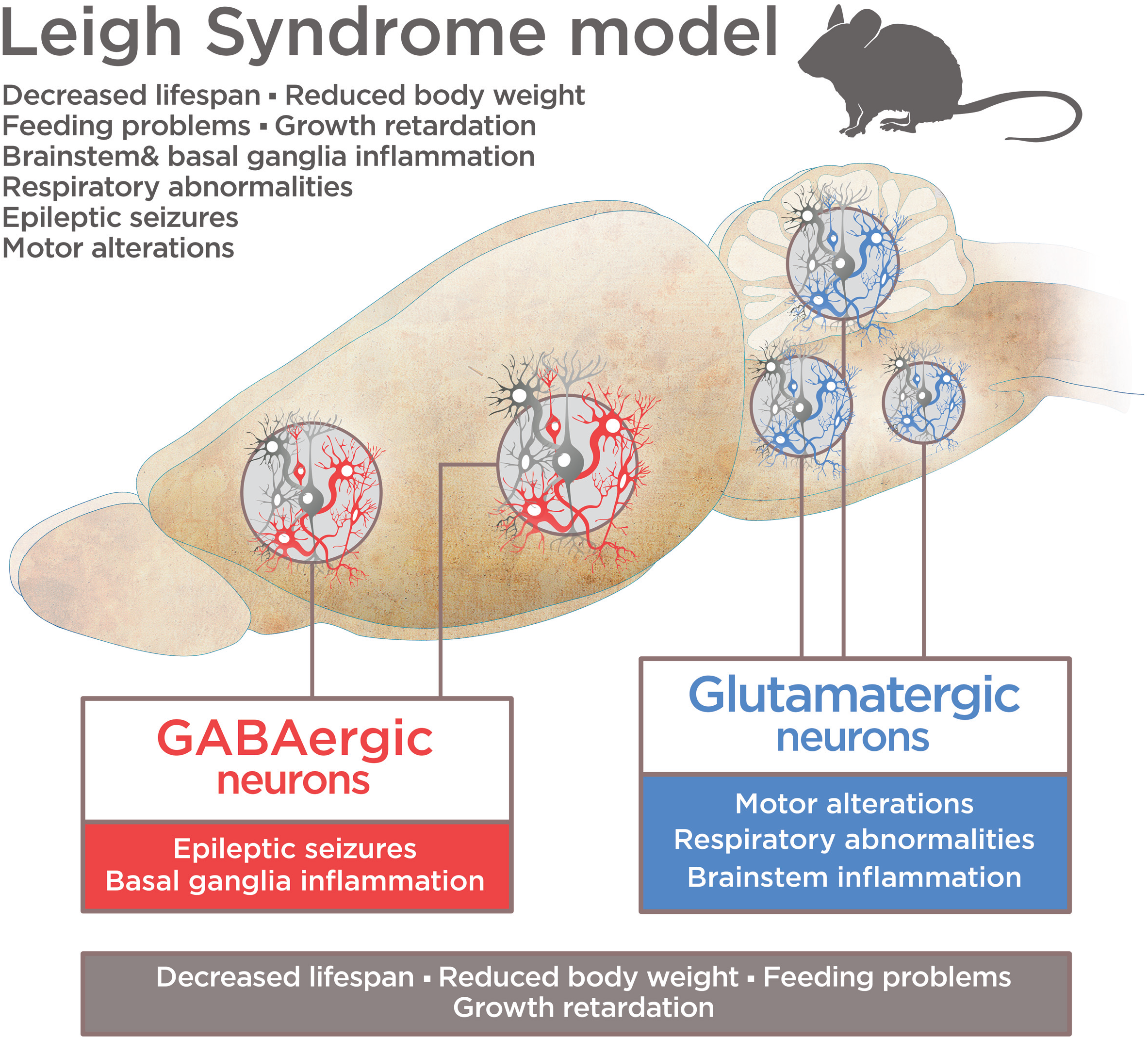
Graphical abstract

In individuals with mitochondrial disease, seizures are commonly observed after stressors such as febrile events^25^. Hence, susceptibility of Gad2:Ndufs4cKO mice to thermally-induced seizures was assessed (Figure 6E-F). Gad2:Ndufs4cKO mice were highly sensitive to temperature compared to Gad2:Ndufs4cCT mice, with some mice displaying seizures already at basal temperature. The number of seizures significantly increased with temperature; 50% of the mice exhibited seizures at 39.5°C and all Gad2:Ndufs4cKO manifested seizures by 41.5°C. Both spontaneous and temperature-induced seizures were always preceded by a myoclonic (MC) seizure (Figure 6A, 6F) with the latter being more severe (Racine scale stage 5) than the former (Racine scale stage 4) (data not shown). Both spontaneous and thermally-evoked seizures were characterized by EEG hyperactivity. Similarly to controls, Vglut2:Ndufs4cKO did not show susceptibility to temperature-induced seizures up to 42°C (data not shown). Daily administration of the anti-epileptic drugs levetiracetam (60 mg/kg) or carbamazepine (40 mg/kg) to Gad2:Ndufs4cKO mice failed to extend their lifespan (Supplemental Figure S7A). Drug-treated mice showed a median survival of 63 and 66 days, respectively, which was not statistically significant when compared to the median lifespan of vehicle-treated mice (59 days). Abnormal activation of the mechanistic target of rapamycin (mTOR) pathway has been involved in epileptic episodes, and its inhibition suppresses seizures in otherwise drug-resistant epilepsy^26–28^. *Ndufs4* deletion leads to elevated mTOR signaling in mice and its inhibition with rapamycin (8 mg/kg) extends lifespan in Ndufs4KO mice^29^. Hence, we administered rapamycin (8 mg/kg; 5 days/week) to Gad2:Ndufs4cKO mice. This treatment significantly inhibited mTOR activity as assessed by the phosphorylation status of its downstream target ribosomal protein S6 (Supplemental Figure S7B-C). However, no effect on survival was observed in rapamycin-treated Gad2:Ndufs4cKO mice compared to vehicle-treated mice (59 days median survival for both groups) (Supplemental Figure S7A).

## Discussion

Neuronal vulnerability is one of the hallmarks of Leigh Syndrome. However, the restricted anatomical distribution of brain lesions indicates a clear gradation in neuronal susceptibility to LS-causing mutations. Specific neuronal populations in affected regions, such as brainstem or basal ganglia, are highly vulnerable to mitochondrial dysfunction and may underlie the plethora of neurological signs observed in LS. The scarcity and high variability of patients has limited our knowledge on the genetic identity and relative contribution of the affected neuronal populations to the phenotype; thus, a model system with consistent neuropathological features resembling mitochondrial disease is a valuable research tool. We have shown^11,13^ that mice lacking the *Ndufs4* gene (Ndufs4KO mice) recapitulate the clinical signs of the human disease^9,13^. Ndufs4KO mice present overt lesions predominantly in the brainstem^11,13,29^, but also in the striatum in late stages of the disease^13^. Hence, we hypothesized that a concerted role of diverse neuronal populations was necessary to drive the plethora of symptoms observed in Ndufs4KO mice.

In this study, we use a conditional genetic approach to selectively ablate NDUFS4 in ChAT-expressing cholinergic neurons (ChAT:Ndufs4cKO), Vglut2-expressing glutamatergic neurons (Vglut2:Ndufs4cKO) or Gad2-expressing GABAergic neurons (Gad2:Ndufs4cKO), thus restricting Complex I deficiency to some of the most abundant neuronal populations. This approach allowed us to provide a comprehensive dissection of the neuronal involvement in the phenotype of LS.

Animals lacking *Ndufs4* in cholinergic neurons (ChAT:Ndufs4cKO) were phenotypically equivalent to controls, indicating no overt contribution of this cell type to the pathology observed in Ndufs4KO mice. In contrast, Vglut2:Ndufs4cKO and Gad2:Ndufs4cKO mice had reduced lifespan and body weight, which was accompanied by a decrease in food intake, which are common clinical signs that appear in Leigh Syndrome patients^1,30^. Recent reports have shown that neuronal cell-specific NDUFS4 knock down in *Drosophila* also leads to severe feeding abnormalities and premature death^31^. Our results indicate a conserved role for neurons in the onset and progression of the pathological condition of global *Ndufs4* deficiency and reveal that both glutamatergic and GABAergic systems contribute to the growth and lethality phenotype.

Apart from the premature death and feeding deficits, Vglut2:Ndufs4cKO and Gad2:Ndufs4cKO mice present two markedly distinct clinical entities (summarized in Table 2). With Vglut2:Ndufs4cKO mice, the lethality was associated with severe motor and respiratory alterations, whereas with Gad2:Ndufs4cKO mice, sudden unexpected death was associated with epilepsy (SUDEP)^32–34^ with no overt clinical alteration beyond the weight loss.

**Table 2.** Clinical signs in *Ndufs4-LS* patients, Ndufs4KO mice and conditional Vglut2:Ndufs4cKO and Gad2:Ndufs4cKO mice.

|  | *Ndufs4*-LS<br>patients | *Ndufs4*KO<br>mouse | *Vglut2:Ndufs4cKO*<br>mouse* | *Gad2:Ndufs4cKO*<br>mouse* |
| --- | --- | --- | --- | --- |
| *Decreased lifespan* | 22/22 | Yes | Yes | Yes |
| *Feeding impairment* | 8/22 | n.d | Yes | Yes |
| *Reduced body weight* | n.a | Yes | Yes | Yes |
| *Brainstem lesions* | 14/22 | Yes | Yes | No |
| *Basal ganglia lesions* | 9/22 | Yes | No | Yes |
| *Ataxia, motor alterations* | 22/22 | Yes | Yes | No |
| *Growth retardation* | 11/22 | Yes | Yes | No |
| *Hypotonia* | 22/22 | Yes | Yes | No |
| *Respiratory abnormalities* | 22/22 | Yes | Yes | No |
| *Increased sensitivity to VAs* | n.a | Yes | Yes** | No** |
| *Seizures* | 4/22 | Yes | No | Yes |

n.d, not determined; n.a, not available. \*Reported in this study. \*\*Reported in *Zimin et al., 2016, Current Biology* 26, 2194–2201. VAs: volatile anesthetics. Adapted from *Quintana et al. J Clin Invest.* 2012;122(7):2359–2368.

Histologically, Vglut2:Ndufs4cKO mice present with prominent neuroinflammation in areas of the brainstem such as the VN, IO and the cerebellar FN, reminiscent of the lesions found in Ndufs4KO mice^11^. We have identified a critical role of brainstem lesions in the development of fatal breathing alterations observed in the Ndufs4KO mice^13^, in agreement with human LS patients^5^. Glutamatergic signaling in the brainstem has been shown to regulate breathing^35^. In addition, VN glutamatergic neurons have been suggested to modulate respiratory responses^36^. Furthermore, the Pre-Bötzinger (PreBotC) complex, a key respiratory center, is composed of glutamatergic neurons^37^ and receives extensive glutamatergic inputs^38^. We have shown that Ndufs4KO mice present intrinsic PreBotC alterations and that vestibular projections to the PreBotC are necessary for maintaining respiration^13^. Our electrophysiological recordings in Vglut2-expressing VN neurons show relatively normal basal firing rate but failure to respond to locomotor activity. Hence, it is likely that failure of glutamatergic VN projections to the PreBotC allowing appropriate responses to physiological needs underlies the breathing deficits observed in Vglut2:Ndufs4cKO mice.

Development of brainstem lesions correlate with motor deficits in animals constitutively lacking *Ndufs4^11^*. Accordingly, strategies that improve motor symptoms in Ndufs4KO mice, such as AAV-based gene therapy^13,39^, rapamycin administration^29^, or hypoxia^40,41^ cause a marked reduction in brainstem lesions. Here, we have identified a critical role for Vglut2-expressing glutamatergic neurons, in the brainstem and cerebellum, in the development of the motor deficits observed after *Ndufs4* deficiency. In keeping with this, conditional deletion of *Ndufs4* in dopaminergic neurons does not cause cell loss^42^ or motor deficits ^43^. However, other areas, such as the striatum, may participate in the delayed, mild, progressive motor dysfunction observed in LS patients^15,39^.

Our gene expression analysis in brainstem of Vglut2:Ndufs4cKO mice has enabled the generation of an in-depth profile of the transcriptomic landscape in an affected brain area after *Ndufs4* deficiency. This analysis revealed a prominent increase in inflammatory mediators in the affected tissue. However, anti-inflammatory or immunotherapeutic approaches have been mostly ineffective as treatments for LS^29,44^ with only a few successful cases reported^45^. Our deconvolution data show a marked infiltration of distinct leucocyte populations, underscoring the complex cellular milieu elicited by mitochondrial dysfunction that may underlie the failure of global anti-inflammatory approaches. Delineation of the immune cells recruited to the brain lesions may lead to novel therapeutic approaches tailored for LS.

The Gad2-expressing GABAergic neurons do not participate in the appearance of respiratory or motor deficits in *Ndufs4* deficiency. However, they are critical for the onset and development of the fatal epileptic seizures that are observed in both global Ndufs4KO mice^11^ and LS patients^21,46^. Lack of *Ndufs4* in Gad2-expressing neurons leads to the appearance of neuroinflammation in the basal ganglia nuclei such as the GPe and SNr. We show that electrophysiological alterations in the GPe neurons predate cortical epileptic events, suggesting a primary role of the basal ganglia circuitry in the development of epileptic seizures in Gad2:Ndufs4cKO mice. Basal ganglia are involved in epilepsy^47–49^, likely by acting as an inhibitory input to cortical seizure spread via feedback mechanisms^47^. Hence, we hypothesize that basal ganglia inhibitory network is affected in Gad2:Ndufs4cKO mice, being unable to control the activity of cortical excitatory neurons, thus leading to epilepsy. Gad2:Ndufs4cKO mice are resistant to different antiepileptic approaches, such as the widely-used antiepileptic drugs carbamazepine and levetiracetam. Epilepsy-induced death in mitochondrial disease patients is usually linked to refractory epileptic seizures^46^. Hence, Gad2:Ndufs4cKO mice may represent an excellent model to study epileptic mechanisms in LS.

Commonly used antiepileptic drugs may promote mitochondrial toxicity, hindering their use in LS^50^. Therefore, non-conventional antiepileptic interventions, such as rapamycin^51^, may represent a better alternative for the treatment of seizures in LS. However, Gad2:Ndufs4KO mice were also resistant to the antiepileptic effects of rapamycin. Previous results have shown that daily rapamycin administration ameliorates the clinical signs and significantly extends lifespan of mice with global *Ndufs4* ablation^29^. While other complications of rapamycin administration, such as immunosuppression and infection may underlie the eventual demise of rapamycin-treated Ndufs4KO mice, our results suggest that rapamycin-resistant phenotypes, such as SUDEP may play a significant role in the mortality of these mice.

In conclusion, we provide new insights on the genetic identity of affected neuronal populations in LS by dissecting the associated cell type-specific molecular, biochemical, clinical and behavioral features in a model of LS. Our work highlights the importance of addressing mitochondrial disease at the cell type-specific level. The advent of new tools to assess transcriptomic and biochemical changes at this level of resolution^52,53^ bodes well for more progress. Hence, our work broadens current understanding of the etiology of LS and paves the way for future studies at the cell type-specific level to unravel the molecular determinants of neuronal pathology in LS.

## Methods

### Study approval

All experiments were conducted following the recommendations in the Guide for the Care and Use of Laboratory Animals and were approved by the Animal Care and Use Committee of the Seattle Children’s Research Institute and the Universitat Autònoma de Barcelona.

### Animal husbandry

Mice were maintained with Teklad Global rodent diet No. 2014S (HSD Teklad Inc., Madison, Wis.) and water available *ad libitum* in a vivarium with a 12-hour light/dark cycle at 22°C.

### Mouse genetics

The following mouse lines were used in this study: *Slc17a6^Cre^* (BAC-Vglut2::Cre)^54^ mice were generated by Ole Kiehn. *Gad2^Cre/+^* (Gad2-IRES-Cre)^55^ and *Chat^Cre/+^* (Chat-IRES-Cre)^56^ mice were obtained from The Jackson Laboratory (Stock No: 028867 and 031661, respectively) (Bar Harbor, ME). *Ndufs4^lox/lox^* and *Ndufs4^Δ/+^* were previously generated by our group^11,12^. Male and female mice of different ages were used in this study. Sex and age of the animals is described in the figure legends. All mice were on a C57BL/6J background after backcrossing for at least 10 generations.

Mice with conditional deletion of *Ndufs4* in Vglut2-expressing glutamatergic neurons *(Slc17a6^Cre^, Ndufs4^Δ/lox^* or Vglut2:Ndufs4cKO) were generated by crossing mice with one *Ndufs4* allele deleted and expressing a codon-improved Cre recombinase (iCre) under the *Slc17a6* promoter (*Slc17a6^Cre^*, *Ndufs4^Δ/+^*) to mice with two floxed *Ndufs4* alleles (*Ndufs4^lox/lox^*). Mice with conditional deletion of *Ndusf4* in Gad2-expressing GABAergic neurons (*Gad2^Cre/+^*, *Ndufs4^lox/lox^* or Gad2:Ndufs4cKO) were obtained by crossing mice with one floxed *Ndufs4* allele and expressing Cre recombinase under the control of the *Gad2* promoter (*Gad2^Cre/+^*, *Ndufs4^lox/+^*) to mice carrying two floxed *Ndufs4* alleles (*Ndufs4^lox/lox^*). Similarly, mice with conditional *Ndufs4* deletion in ChAT-expressing cholinergic neurons (*Chat^Cre/+^*, *Ndufs4^lox/lox^* or ChAT:Ndufs4cKO) were obtained by crossing mice carrying one floxed *Ndufs4* allele and expressing Cre recombinase driven by the ChAT promoter (*Chat^Cre/+^*, *Ndufs4^lox/+^* mice) to *Ndufs4^lox/lox^* mice. Littermate controls were *Slc17a6^Cre^*, *Ndufs4^lox/+^* (Vglut2:Ndufs4cCT); *Gad2^Cre/+^*, *Ndufs4^lox/+^* (Gad2:Ndufs4cCT) and *Chat^Cre/+^* *Ndufs4^lox/+^* (ChAT:Ndufs4cCT) mice. In all cases, genotype of the offspring was determined by PCR analysis. Primer sequences have been described^12^.

### Clinical evaluation

Vglut2:Ndufs4cKO and Gad2:Ndufs4cKO mice were examined every other day for clinical signs resulting from cell type-specific *Ndufs4* inactivation. Physiological (body weight) and behavioral (locomotor activity, motor coordination, gait/postural alterations) parameters were evaluated in more than 50 animals for each mouse line and were grouped into the following categories based on visual observation: “+++” severe manifestation of the clinical sign, “++” moderate manifestation of the clinical sign, “+” mild clinical sign, “-” absence of clinical sign. Mice were humanely euthanized after losing 20% of their peak body weight.

### Food intake analysis

Food consumption was recorded from 7 to 11 weeks of age using a Physiocage system (Panlab, Spain). Data at 8 weeks of age (right before the median survival value) is presented, including enough individuals to ensure sufficient statistical power.

### Behavioral assays

#### Rotarod test

A standard rotarod device (Rotarod, San Diego Instruments, USA) was used to assess motor coordination and global physical condition of animals. Mice were placed on the spindle, which linearly accelerated from 4 to 40 r.p.m and increased 2 r.p.m. every 10 seconds. Each mouse received five trials per day over 3 days with a 5 min rest period between trials. The trial ended when the mouse fell off the spindle or after 3 minutes (cutoff time).

#### Open field

Mice were placed in the open field arena (560 [W] × 365 [D] × 400 [H] mm) and allowed to move freely for 10 min. Locomotor activity of mice was next monitored and total distance traveled (m) and velocity (cm/s) measured using the EthoVision tracking software (Noldus).

#### Whole–body plethysmography

Ventilatory function was assessed by whole-body plethysmography (EMMS, England, UK) under unrestrained conditions. The system was calibrated to 1 ml volume. An acclimation period of 45 min was allowed for mice adaptation to the chamber, followed by a 15-min experimental period. For Vglut2:Ndufs4cKO mice, plethysmography recordings were performed at different stages of the disease (early, n=5; mid, n=7 and late, n=12) according to the clinical examination, and compared to littermate controls (n=12). Studies were also conducted on Gad2:Ndufs4cKO mice (n=7) between 50-60 days of age and compared to corresponding controls (n=7). Respiratory frequency (F_R_; breaths-minute^-1^) and tidal volume normalized per body weight (V_T_; μL-g^-1^) were measured.

#### Tissue preparation

For immunofluorescence, mouse brains were collected and fixed overnight in 4% paraformaldehyde (PFA) in PBS (pH 7.4). Subsequently, brains were cryoprotected in a PBS solution containing 30% sucrose and frozen in dry ice. Frozen brains were embedded in OCT, sectioned at 30 μm in a cryostat and rinsed in PBS prior to staining. For western blot analysis, brain areas (olfactory bulb, thalamus, spinal cord and globus pallidus) were dissected according to the Paxinos mouse brain atlas^57^ and flash-frozen in liquid nitrogen before homogenization.

#### Immunofluorescence

Tissue sections were rinsed in PBS-0.2% Triton X-100 (PBST) solution. Non-specific binding was blocked with 10% normal donkey serum (NDS) in a PBST solution for 60 min at room temperature, followed by overnight incubation at 4°C with primary antibodies diluted in 1% NDS-PBST (1:2,000 for mouse anti-GFAP, Sigma; 1:1,000 for chicken anti-GFAP, Abcam; 1:1,000 for anti-Iba-1, Wako; 1:1,000 for anti-TH, Millipore). Sections were then washed in PBST and incubated for 1h at room temperature with the corresponding Cy-(1:200, Jackson Immunoresearch) or Alexa Fluor-conjugated secondary antibodies (1:500, Thermo Fisher Scientific) in 1% NDS-PBST. Sections were finally washed in PBS and rinsed in water before mounting onto slides with Fluoromount G (Electron Microscopy Sciences) for microscopic analysis.

#### Western Blotting

Brain tissue samples were homogenized in iced-cold RIPA buffer (Santa Cruz Biotechnology) and protein concentration determined by the BCA assay (Thermo Fisher Scientific). Thereafter, 20 μg of protein lysates were heat-denatured in Laemmli sample buffer (Bio-Rad Laboratories, Inc.), subjected to 4-20% gradient SDS-PAGE and transferred to nitrocellulose membranes (Bio-Rad Laboratories, Inc.). Membranes were then blocked for 1 hour with 5% (w/v) dried skimmed milk in Tris-buffered saline containing 0.1% Tween-20 (TBS-T) and incubated overnight at 4°C with primary antibodies against NDUFS4 (Abcam, mouse, 1:500), NSE (Dako, mouse, 1:1,000), GFAP (Sigma, mouse, 1:50,000), Iba1 (Wako, rabbit, 1:10,000), p-S6(Ser235/236) (Cell signaling, rabbit, 1:2,000), β-actin (Sigma, mouse, 1:20,000) or GAPDH (GeneTex, mouse, 1:40,000). After incubation with the corresponding HRP-conjugated secondary antibodies (1:10,000; Jackson ImmunoResearch), membranes were washed in TBS-T and developed using an enhanced chemiluminescence (ECL) detection system (Pierce). Bands were quantified using Image J software (National Institutes of Health, USA).

#### Whole-genome gene expression (WGGEX) analysis

For WGGEX analysis, 150 ng of total RNA extracted from the brainstem of Vglut2:Ndufs4cKO (n=4) and Vglut2:Ndufs4cCT mice (n=4) was amplified and biotin-labeled using the Illumina TotalPrep RNA Amplification kit (Ambion). 750 ng of the labeled cRNA was hybridized to MouseRef-8 v2 expression beadchips (Illumina) for 16 h before washing and analyzing according to the manufacturer’s directions. Signal was detected using a BeadArray Reader (Illumina), and data were analyzed for differential expression using the GenomeStudio data analysis software (Illumina). Average normalization, the Illumina custom error model, and multiple testing corrections using the Benjamini and Hochberg false discovery rate were applied to the analysis. Only transcripts with a differential score of >13 *(P* < 0.05) were considered. Normalized and raw data have been deposited in the National Center for Biotechnology Information Gene Expression Omnibus (accession number GSE125470). Functional enrichment analysis of differentially expressed mRNAs (1.4 fold or higher) using overrepresentation analysis (ORA) was accomplished using WebGestalt http://www.webgestalt.org)^16^. Characterization of the immune cell composition in these gene expression profiles was accomplished using the computational algorithm ImmuCC^18^.

#### Surgery

Mice underwent survival surgery to implant EEG and EMG electrodes under isoflurane anesthesia with subcutaneous bupivacaine (1 mg/kg) for analgesia as described^58^. Using aseptic technique, a midline incision was made anterior to posterior to expose the cranium. Each EEG electrode consisted of a micro-screw attached to a fine diameter (130 μm bare; 180 μm coated) silver wire. The screw electrodes were placed through the small cranial burr holes at visually identified locations: Left and right frontal cortices approximately 1 mm anterior to the bregma and 3 mm lateral of the sagittal suture. EMG electrodes were placed in back muscles. A reference electrode was placed at the midline cerebellum and a ground electrode was placed subcutaneously over the back and the skin was closed with sutures. All electrodes connected to a micro-connector system and their impedances were typically < 10 kΩ. After electrode placement, the skin was closed with sutures and the mice were allowed to recover from surgery for 2-3 days.

#### Video-EEG-EMG

Recording approach was performed as described^58^. Simultaneous video-EEG-EMG records were collected in conscious mice on a PowerLab 8/35 data acquisition unit using Labchart 7.3.3 software (AD Instruments, Colorado Spring, Co). All bio-electrical signals were acquired at 1 KHz sampling rate. The EEG signals were processed off-line with a 1-80 Hz bandpass filter and the EMG signals with a 3-Hz highpass filter. Video-EEG-EMG data collected were analyzed using Labchart software.

#### Thermal seizure induction

Mouse body temperature was controlled using a rectal temperature probe and a heat lamp attached to a temperature controller in a feedback loop (Physitemp Instruments Inc., NJ). Briefly, as described^59^, body temperature was increased by 0.5°C every 2 minutes until seizure occurred or a 42°C temperature was reached. Mice were immediately cooled using a small fan.

### Electrophysiological studies

#### Surgery

Vglut2:Ndufs4cKO and Gad2:Ndufs4cKO mice and their respective controls (30 to 40 days old) were anesthetized with 1.5% isoflurane and implanted with a homemade implant in either the vestibular nuclei (AP: −6.0; ML: −1.0; DV: −4.00) or the lateral globus pallidus (AP: −0.46; ML: −1.95; DV: −4.00), according to Paxinos^57^. Briefly, a small craniotomy window was made above the desired recording site and a 4-tetrode bundle was lowered into the brain until its destination at 0.1 mm/sec (Robot Stereotaxic, Neurostar, Germany). As ground, a stainless steel wire (0.075 mm diameter, Advent Research Material Ltd., England) was placed between skull and dura over the cerebellum or visual cortex. Then the entire implant was attached and secured to animal head with dental cement. After surgery, animal welfare and body weight were documented daily and mice allowed to recover for 10-14 days. In Vglut2:Ndufs4cKO mice and their respective controls, a viral vector expressing the light-sensitive cation channel Channelrodopsin-2 in a Cre-dependent manner (AAV-DIO-ChR2) was delivered into the VN prior to implantation of the tetrode to identify glutamatergic Vglut2-expressing cells after blue light (473 nm) stimulation.

#### Electrophysiological data acquisition

To explore presence of seizures, the electrophysiological activity was recorded from Gad2:Ndufs4cCT and Gad2:Ndufs4cKO freely moving mice. After the recovery period, animals were placed on a daily basis in an open field arena and both local field potentials (LFPs) and extracellular single-unit activity were recorded until the death of the animal. In Vglut2:Ndufs4cCT and Vglut2:Ndufs4cKO mice, a tetrode bundle was attached along a fiber optic (Ø200 μm Core, 0.22 NA, FG200UCC, Torlabs, USA) to deliver light and record neuronal cells at the same time. Again, all animals were daily recorded in an open field until death.

For all experiments, local field potential activity was amplified, A-D converted and sampled at 1KHz and bandpass filtered at 0.1 to 250 Hz (DigitalLynx SX and Cheetah Data Acquisition System, Neuralynx, USA). Continuous spike signals were also recorded, amplified, band-pass filtered (300 Hz to 8 kHz) and sampled at 32 kHz. Optogenetic stimulations (ranging from 50 to 200-ms blue light stimuli, 473 nm, delivered frequency from 1 to 40 Hz) were delivered through a blue-emitting diode laser (473nm DPSS Laser System, Laserglow Technologies, CA).

#### Electrophysiological data analysis

For the presence of seizures, data were analyzed in Spike2 (Cambridge Electronic Design Limited, UK) for visual inspection and report presence of epileptic events. Offline single-unit spike sorting was performed with Offline Sorter software (Offline Sorter, Plexon Inc., USA). Briefly, for each channel, a specific manual threshold was defined and all events bypassing this threshold were assumed to be an action potential. After an overall waveform shape and 3 principal component analysis (PCA) inspection, all spikes were sorted with an automatic K-mean algorithm to separate clusters of cells. After sorting, a final inspection of ISI histogram and sorting statistics (MANOVA F statistics, J3 and the Davies-Bouldin validity index) was performed to ensure the best single-unit clustering. In Vglut2:Ndufs4cCT and Vglut2:Ndufs4cKO mice, to identify glutamatergic neurons from all sorted cells, response to optogenetic stimulations were plotted. For all stimulation trains, during a baseline activity corresponding to 500ms before stimulation, the mean number of spikes per bins was calculated. Then, a single cell was identified as glutamatergic only if, during optogenetic stimulation, there were bins expressing a number of spikes superior to the baseline mean plus 3 times the standard deviation of the baseline mean. Finally, after analysis of spiking activity (Matlab, MathWorks, USA), cells with a coefficient of variation of interspike interval (CV.ISI) over 3 were exclude from the data set.

#### Drug treatments

Levetiracetam (Keppra^®^ parental formulation), rapamycin (MedChemExpress) and carbamazepine (Sigma-Aldrich) antiepileptic drugs were injected i.p. in Gad2:Ndufs4cKO mice starting at postnatal day 40. Levetiracetam (6 mg/mL in saline solution) was administrated i.p. at 60 mg/kg every day (n=3). A stock solution of rapamycin (50 mg/mL) was prepared in absolute ethanol and stored at −20°C. Prior to injections, this was further diluted to 0.4 mg/mL in saline with 4% ethanol, 5% polyethylene glycol (PEG) 300 and 5% Tween 80. Rapamycin was injected five days per week at 8 mg/kg (n=3). Carbamazepine was first diluted to 20 mg/mL in PEG 300, and further diluted to 8 mg/mL in saline for daily administration (i.p.) at 40 mg/kg (n=5). Control mice were injected with respective vehicles.

#### Statistics

Data are shown as the mean ± SEM. GraphPad Prism v5.0 software was used for statistical analyses. Appropriate tests were selected depending on the experimental design as stated in the figure legends. Statistical significance, when reached (*P* <0.05 was considered significant), is stated in the figure legends.

## Author contributions

IB, AG, ES, PPD, FM, PM, FK, AQ conducted experiments and acquired and analyzed the data. IB, AG, ES, FK and AQ designed research studies and wrote the manuscript. AQ and FK provided reagents and AQ coordinated the work.

## Acknowledgments

This work was supported by a Marie Sklodowska-Curie Individual Fellowship (H2020-MSCA-IF-2014-658352; ES), a Marie Sklodowska-Curie COFUND action (H2020-MSCA-COFUND-2014-665919; AG), a Juan de la Cierva fellowship (JCI-2015-24576; IB), a pre-doctoral fellowship (BES-2015-073041; PPD) and a Ramón y Cajal fellowship (RyC-2012-11873; AQ). A.Q. received funds from the Seattle Children’s Research Institute and Northwest Mitochondrial Guild, and grants from the European Research Council (Starting grant NEUROMITO, ERC-2014-StG-638106), MINECO Proyectos I+D de Excelencia (SAF2014-57981P; SAF2017-88108-R) and AGAUR (2017SGR-323). We thank Richard Palmiter, Diane Durnam and members of the Quintana lab for insightful discussions, comments and edits.

