## Supplemental Figures for "Defined neuronal populations drive fatal phenotype in Leigh Syndrome"

#### Supplemental Figure legends and Figures:

**Supplemental Figure S1. Generation and characterization of a conditional mouse line lacking *Ndufs4* in ChAT-expressing neurons.**

(A) Breeding strategy used to generate mice with selective *Ndufs4* deletion in cholinergic neurons (ChAT:Ndufs4cKO mice) and its genetic control (ChAT:Ndufs4cCT mice). Body weight (B) and rotarod performance (C) in ChAT:Ndufs4cKO mice and their respective control (ChAT:Ndufs4cCT). Data are presented as the mean ± SEM. Statistical analysis was performed using a two-way ANOVA.

**Supplemental Figure S2. NDUFS4 levels and food intake after conditional deletion of *Ndufs4* in Vglut2- and Gad2-expressing cells.** (A) Western blot analysis for

NDUFS4 and GAPDH (loading control) levels in the olfactory bulb and thalamus of Vglut2:Ndufs4cKO mice, Gad2:Ndufs4cKO mice, and their respective controls (Vglut2:Ndufs4cCT and Gad2:Ndufs4cCT mice) (n=3). (B) Food intake (Kcal/day) corrected for body weight (g) in Vglut2:Ndufs4cKO mice (n=6), Gad2:Ndufs4cKO mice (n=5), and their respective controls (Vglut2:Ndufs4cCT, n=7, and Gad2:Ndufs4cCT, n=4). Data are presented as the mean ± SEM. Statistical analysis was performed using an unpaired *t*-Test.

**Supplemental Figure S3. Reactive gliosis in the spinal cord of Vglut2:Ndufs4cKO mice.** (A) Vglut2:Ndufs4cKO mice present hindlimb clasping starting at P40 (B) Western blot analysis (top) and band intensity quantification (bottom) showing GFAP, Iba-1, NSE and GAPDH levels in spinal cords of Vglut2:Ndufs4cCT and Vglut2:Ndufs4cKO mice (n=5). GAPDH was used as a loading control. Data are shown as mean ± SEM. Statistical analysis was performed using an unpaired *t*-Test $^{**}P < 0.01$, $^{***}P < 0.001$. (C) Immunofluorescence analysis for GFAP and Iba-1 (top), hematoxylin and eosin staining (H&E; middle), and Luxol fast blue staining (LFB; bottom) in spinal cord sections from Vglut2:Ndufs4cKO and Vglut2:Ndufs4cCT mice.

**Supplemental Figure S4. *Ndufs4* deficiency in Gad2-expressing GABAergic and Vglut2-expressing glutamatergic cells results in prominent neuroinflammation in specific areas.** (A) Immunofluorescence analysis for tyrosine hydroxylase (TH; dopaminergic neurons), GFAP (astrocytes) and Iba-1(microglia) in brain sections of Gad2:Ndufs4cCT and Gad2:Ndufs4cKO mice containing the substantia nigra (SN). (B)

Immunofluorescence analysis for Calbindin (Calb; Purkinje cells) and Iba-1 in brain sections of Gad2:Ndufs4cKO mice containing the cerebellar vermis (top) and the cerebellar flocculus (bottom). DAPI was used as a nuclear counterstain. (C) Volcano plot showing differentially expressed transcripts ($P<0.05$, Fold Change (FC) >2) in the brainstem of Vglut2:Ndufs4cKO mice when compared to Vglut2:Ndufs4cCT mice.

**Supplemental Figure S5. Age-dependent motor decline in Vglut2:Ndufs4cKO mice.** Total distance travelled (A) and speed (B) was analyzed in the rotarod test for Vglut2:Ndufs4cKO and Vglut2:Ndufs4cCT at different weeks of age (6-9 weeks). Data are presented as the mean ± SEM. Statistical analysis was performed using two-way ANOVA with Bonferroni post-test (*P < 0.05; ***P < 0.001).

**Supplemental Figure S6. Vglut2:Ndufs4cKO mice do not present spontaneous seizures.** EEG-EMG recordings (A-B; B is a magnification of the inset in A) and spectrogram analysis (C) in a Vglut2:Ndufs4cKO mouse.

**Supplemental Figure S7. Lifespan of Gad2:Ndufs4cKO mice is not extended by antiepileptic treatment.** (A) Survival rate curve for Gad2:Ndufs4cKO mice treated intraperitoneally (i.p) with vehicle (black line), levetiracetam (60 mg/kg daily, n=3, red line), rapamycin (8 mg/kg; 5 days/week, n=3, blue line) or carbamazepine (40 mg/kg daily, n=5, green line). Western blot analysis (B) and quantification (C) for p-S6(Ser235/236) and β-actin (loading control) levels in lysates from the globus pallidus of Gad2:Ndufs4cKO mice treated with 8 mg/kg of rapamycin (n=3) or vehicle (n=4). Data are shown as mean ± SEM. Statistical analysis was performed using an unpaired *t*-Test **P < 0.01.

**Supplemental Video S1**. Breathing irregularities in Vglut2:Ndufs4cKO mice.

**Supplemental Video S2**. Generalized tonic clinic seizure without hyper motor (Racine scale stage 4).

### Figure S1

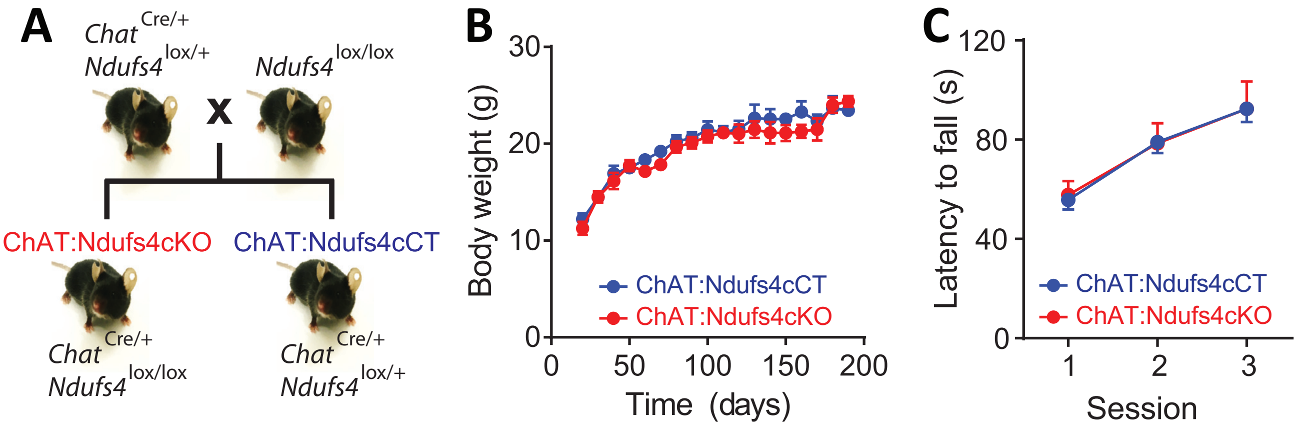
A
Chat Cre/+
Ndufs4 lox/+
x
Ndufs4 lox/lox
ChAT:Ndufs4cKO
ChAT:Ndufs4cCT
Chat Cre/+
Ndufs4 lox/lox
Chat Cre/+
Ndufs4 lox/+
B
Body weight (g)
30
20
10
0
0
50
100
150
200
Time (days)
ChAT:Ndufs4cCT
ChAT:Ndufs4cKO
C
Latency to fall (s)
120
80
40
0
1
2
3
Session
ChAT:Ndufs4cCT
ChAT:Ndufs4cKO

### Figure S2

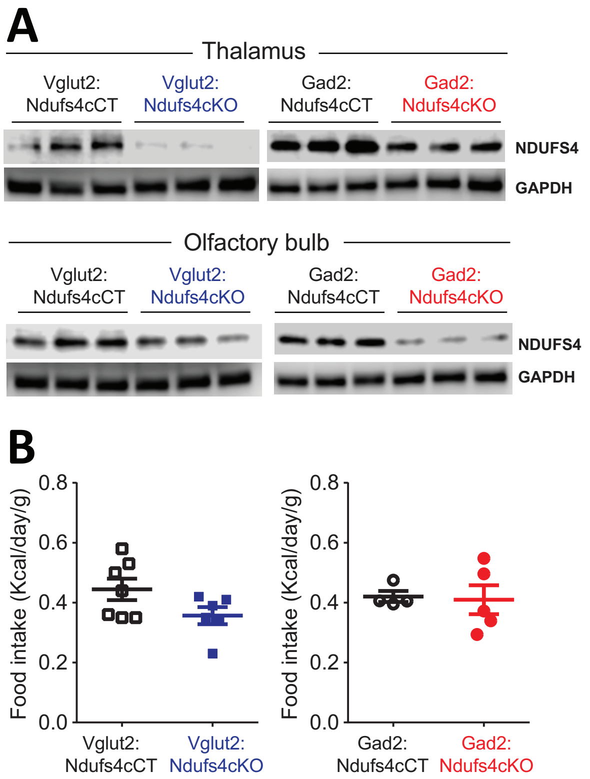
A
Thalamus
Vglut2: Ndufs4cCT
Vglut2: Ndufs4cKO
Gad2: Ndufs4cCT
Gad2: Ndufs4cKO
NDUFS4
GAPDH
Olfactory bulb
Vglut2: Ndufs4cCT
Vglut2: Ndufs4cKO
Gad2: Ndufs4cCT
Gad2: Ndufs4cKO
NDUFS4
GAPDH
B
Food intake (Kcal/day/g)
0.8
0.6
0.4
0.2
0.0
Vglut2: Ndufs4cCT
Vglut2: Ndufs4cKO
Food intake (Kcal/day/g)
0.8
0.6
0.4
0.2
0.0
Gad2: Ndufs4cCT
Gad2: Ndufs4cKO

### Figure S3

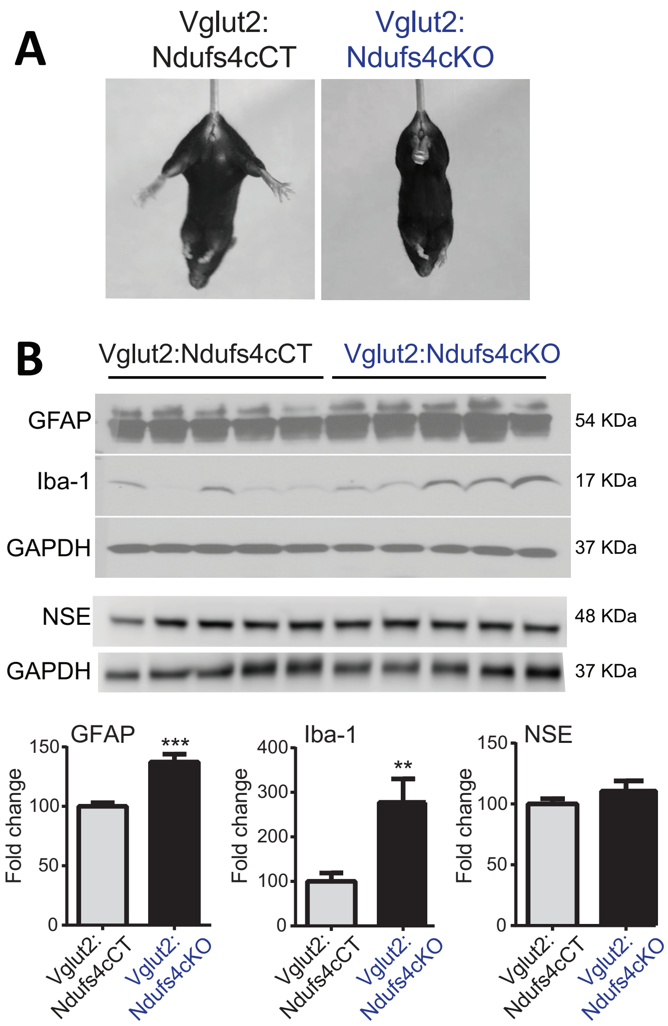

A
Vglut2: Ndufs4cCT
Vglut2: Ndufs4cKO
B
Vglut2:Ndufs4cCT
Vglut2:Ndufs4cKO
GFAP
54 KDa
Iba-1
17 KDa
GAPDH
37 KDa
NSE
48 KDa
GAPDH
37 KDa
GFAP
***
Fold change
150
100
50
0
Vglut2: Ndufs4cCT
Vglut2: Ndufs4cKO
Iba-1
**
400
300
200
100
0
NSE
C
Vglut2:Ndufs4cCT
Vglut2:Ndufs4cKO
GFAP Iba1 DAPI
H & E
LFB

### Figure S4

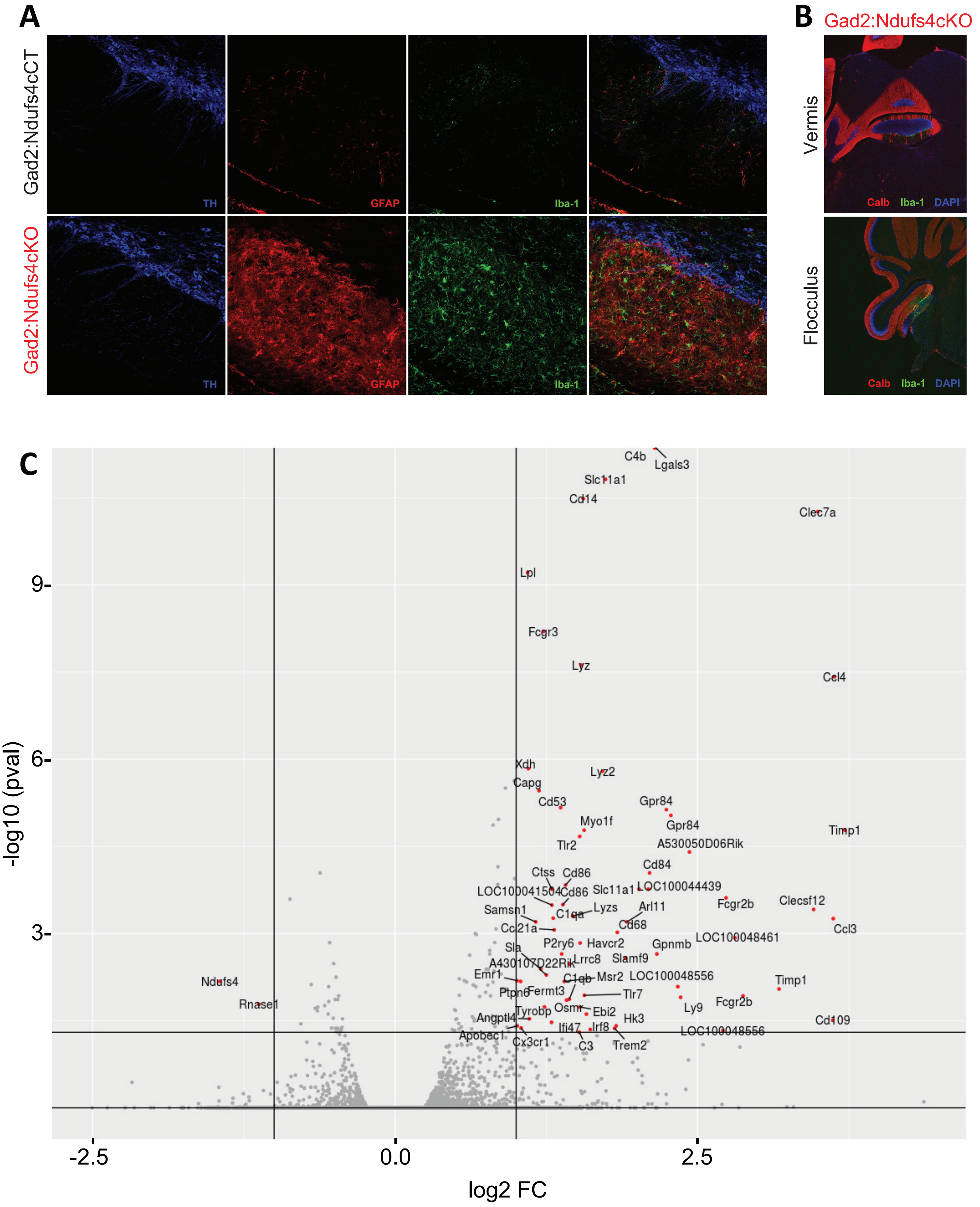
A
Gad2:Ndufs4cCT
Gad2:Ndufs4cKO
TH
GFAP
Iba-1
B
Gad2:Ndufs4cKO
Vermis
Flocculus
Calb
Iba-1
DAPI
C
C4b
Lgals3
Slc11a1
Cd14
Clec7a
Lpl
Fcgr3
Lyz
Ccl4
Xdh
Lyz2
Capg
Cd53
Gpr84
Myo1f
Gpr84
Timp1
Tlr2
A530050D06Rik
Cd84
Ctss
Cd86
LOC100041504
Cd86
Slc11a1
LOC100044439
Fcgr2b
Clecsf12
Samsn1
C1qa
Lyzs
Arl11
Ccl21a
Cd68
Ccl3
P2ry6
Havcr2
Gpnmb
LOC100048461
Sla
A430107D22Rik
Lrrc8
Slamf9
Emr1
C1qb
Msr2
LOC100048556
Timp1
Ndufs4
Ptpn6
Fermt3
Tlr7
Rnase1
Osmr
Ebi2
Ly9
Fcgr2b
Tyrobp
Angptl4
Hk3
Cd109
Ifi47
Irf8
Apobec1
LOC100048556
Cx3cr1
C3
Trem2
9-
6-
3-
-2.5
0.0
2.5
-log10 (pval)
log2 FC

### Figure S5

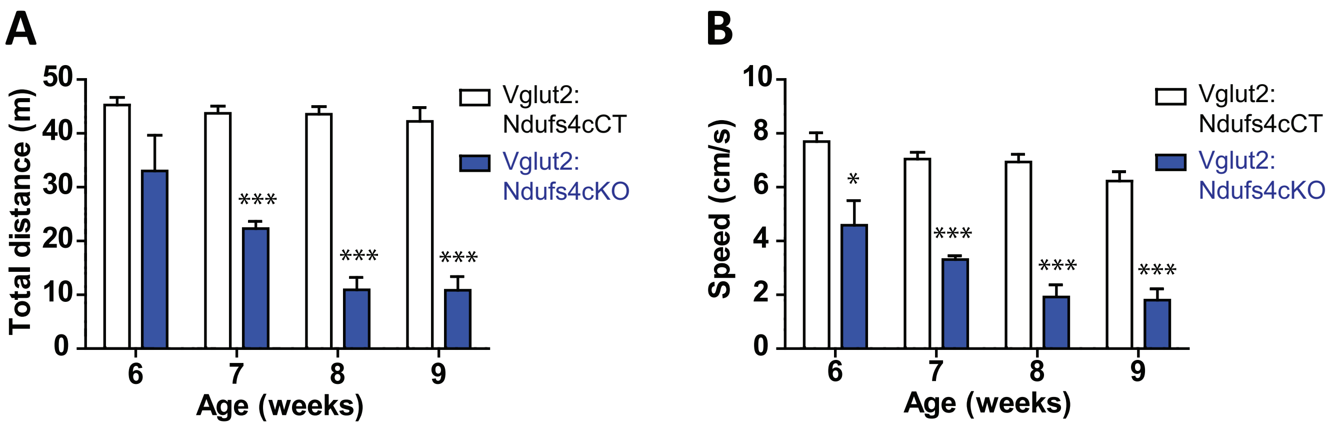
A
Total distance (m)
50
40
30
20
10
0
***
***
***
6
7
8
9
Age (weeks)
Vglut2:
Ndufs4cCT
Vglut2:
Ndufs4cKO
B
Speed (cm/s)
10
8
6
4
2
0
*
***
***
***
6
7
8
9
Age (weeks)
Vglut2:
Ndufs4cCT
Vglut2:
Ndufs4cKO

### Figure S6

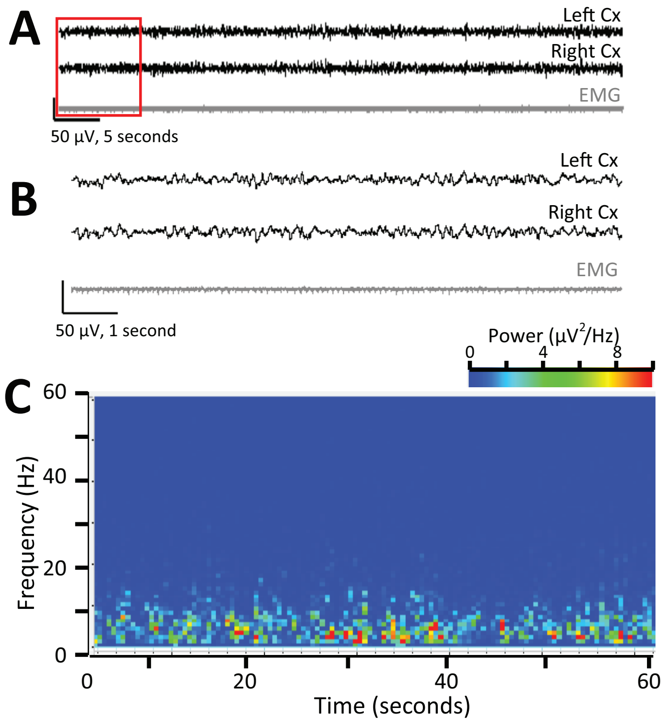
A
Left Cx
Right Cx
EMG
50 µV, 5 seconds
B
Left Cx
Right Cx
EMG
50 µV, 1 second
C
Power (µV²/Hz)
0
4
8
60
40
20
0
Frequency (Hz)
0
20
40
60
Time (seconds)

### Figure S7

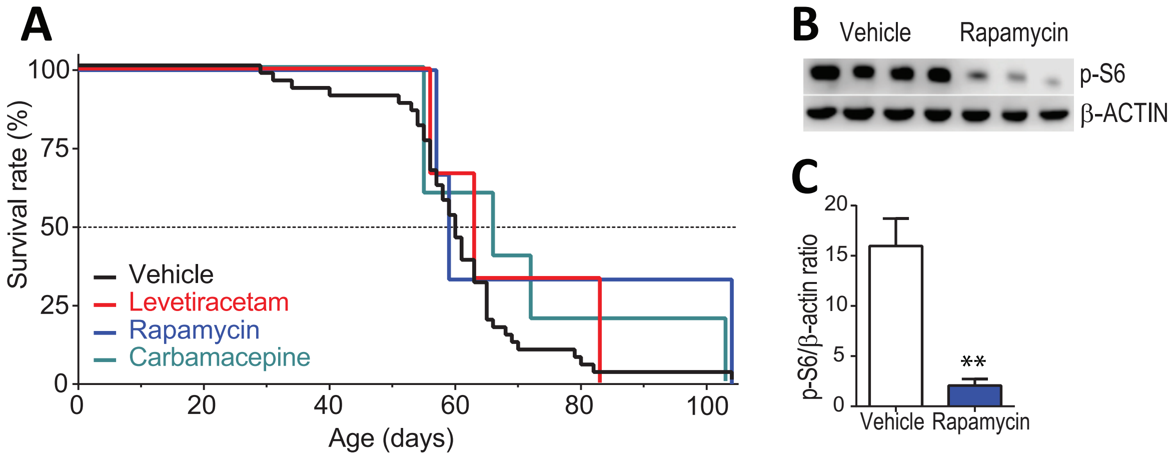
A
Survival rate (%)
100
75
50
25
0
0
20
40
60
80
100
Age (days)
Vehicle
Levetiracetam
Rapamycin
Carbamacepine
B
Vehicle
Rapamycin
p-S6
β-ACTIN
C
p-S6/β-actin ratio
20
15
10
5
0
**
Vehicle
Rapamycin
